# Alcohol Effects on Globus Pallidus Connectivity: Role of Impulsivity and Binge Drinking

**DOI:** 10.1101/819458

**Authors:** Samantha J. Fede, Karina P. Abrahao, Carlos R. Cortes, Erica N. Grodin, Melanie L. Schwandt, David T. George, Nancy Diazgranados, Vijay A. Ramchandani, David M. Lovinger, Reza Momenan

## Abstract

Despite the harm caused by binge drinking, the neural mechanisms leading to risky and disinhibited intoxication-related behaviors are not well understood. Evidence suggests that the globus pallidus externus (GPe), a substructure within the basal ganglia, participates in inhibitory control processes, as examined in stop-signaling tasks. In fact, studies in rodents have revealed that alcohol can change GPe activity by decreasing neuronal firing rates, suggesting that the GPe may have a central role in explaining impulsive behaviors and failures of inhibition that occur during binge drinking. In this study, twenty-five healthy volunteers underwent intravenous alcohol infusion to achieve a blood alcohol level of 0.08 g/dl, which is equivalent to a binge drinking episode. A resting state functional magnetic resonance imaging scan was collected prior to the infusion and at binge-level exposure. Functional connectivity analysis was used to investigate the association between alcohol-induced changes in GPe connectivity, drinking behaviors, and impulsivity traits. We found that individuals with greater number of drinks or heavy drinking days in the recent past had greater alcohol-induced deficits in GPe connectivity, particularly to the striatum. Our data also indicated an association between impulsivity and alcohol-induced deficits in GPe – frontal/precentral connectivity. Moreover, alcohol induced changes in GPe-amygdala circuitry suggested greater vulnerabilities to stress-related drinking in some individuals. Taken together, these findings suggest that alcohol may interact with impulsive personality traits and drinking patterns to drive alterations in GPe circuitry associated with behavioral inhibition, possibly indicating a neural mechanism by which binge drinking could lead to impulsive behaviors.

## Introduction

Harmful drinking behaviors like binge drinking contribute to 5.9% of all global deaths [1, 2]. This is largely due to organ damage caused by chronic drinking, intoxication-related car accidents and domestic violence, and the development of alcohol use disorder (AUD) [3, 4]. Binge drinking is defined as drinking that results in blood alcohol concentration (BAC) levels at or above 0.08 g/dl (estimated as consuming at least 4 or 5 standard drinks in less than two hours for women and men, respectively). The mechanisms underlying binge drinking and related risky behaviors are unclear, but impulsivity or failure in inhibitory control during decision-making likely plays a role [5-8].

Research into the neural correlates of failure to inhibit suggests that the basal ganglia plays an important role [9, 10]. The basal ganglia are formed by several interconnected brain regions that control action selection, reward, goal-directed behavior and habitual learning [11]. Frontostriatal pathways enable inhibitory control quickly via a hyperdirect pathway through the inferior frontal gyrus (IFG) and presupplementary motor area (preSMA) that quickly “brakes”, and through indirect pathways that “suppresses” movement [12, 13]. In particular, the globus pallidus externus (GPe), a central and highly interconnected component of the basal ganglia, may have a key role in inhibitory control [14]. A specific projection from the GPe to the striatum known as the arkypallidal projection has been shown to inhibit behavior temporarily in “stop-signal”-based tasks [15, 16]. Given this role, alterations to GPe connectivity could contribute to increased impulsivity and failure of inhibitory control.

Alcohol has been shown to have a direct effect on GPe neurons. In a mouse model, Abrahao et al. [17] have demonstrated that acute alcohol decreases the firing rate of specific neurons in the GPe, including (1) the low frequency prototypical neurons, which project downstream to the subthalamic nucleus / substantia nigra and upstream to the striatum, and (2) the arkypallidal neurons, which project to the striatum. In humans, alcohol decreased functional magnetic resonance imaging (fMRI) signal in the GP during a stop-signal task [18], and this signal change was associated with slowed reaction time and impaired inhibitory control [10]. Taken together, these findings suggest that impulsivity and behavioral disinhibition associated with alcohol consumption may be driven by alcohol-induced changes in GPe connectivity.

In this study, we aimed to determine whether alcohol-induced changes in GPe connectivity are related to recent drinking patterns, impulsivity, and interactions between the two. To do so, we compared the resting-state fMRI connectivity of the GPe before and after controlled acute intravenous (IV) alcohol administration to “binge drinking” levels (BAC = 0.08 g/dl) in healthy humans, and examined the relationship between those differences, recent drinking history, and trait impulsivity. To our knowledge, this is the first time such an investigation has been conducted. We hypothesized that alcohol would result in greater decreases in GPe functional connectivity across the brain in individuals with more impulsive traits and higher risk drinking behaviors. Moreover, we hypothesized that the interaction between recent drinking behaviors and impulsivity would contribute to alcohol-induced changes in GPe functional connectivity.

## Methods

### Participants

Twenty-five healthy (13 males, 12 females; mean age: 28.85 years), light social drinkers were recruited by local advertisement, according to approved National Institutes of Health Institutional Review Board procedures. The demographic data of participants is summarized in Table 1. After obtaining written informed consent, all participants underwent a comprehensive medical screen, including blood work, urinalysis, medical history, physical exam, and a Structured Clinical Interview for the DSM Axis-I Disorders (SCID-IV) [19]. Criteria for inclusion in this study were: healthy, 21-45 years old, consumption of 1 to 10 drinks per week for females and 1 to 14 drinks per week for males. Subjects also had to have consumed at least two standard drinks of alcohol within one hour on at least one occasion in their lifetime. Criteria for exclusion included: abnormal blood or urine lab test values or findings from the medical screen, DSM-IV criteria for alcohol or other substance dependence (excluding nicotine) at any time; current or past major psychiatric disorder (DSM-IV Axis I), head injury requiring hospitalization, Body Mass Index (BMI) value over 30, inability to stop taking any medication or drugs 3 days prior to study days, and MRI contraindications. Also, non-drinkers were excluded from the study due to ethical concerns related to alcohol administration.

**Table 1.** Demographic, Drinking, and Impulsivity Characteristics

| Variable | Count |  |
| --- | --- | --- |
| Total n | 25 |  |
| Gender (Male / Female) | 13 / 12 |  |
| Smokers | 1 |  |
|  | Mean | SD |
| Age | 28.85 | 7.40 |
| Years Of Education | 15.88 | 1.88 |
| BIS: Total | 54.76 | 9.62 |
| Attentional Impulsiveness | 12.96 | 2.99 |
| Motor Impulsiveness | 20.92 | 3.32 |
| Nonplanning Impulsiveness | 19.72 | 4.21 |
| UPPS-P: Positive Urgency | 1.36 | 0.31 |
| Negative Urgency | 1.66 | 0.44 |
| Premeditation | 1.78 | 0.42 |
| Perseverance | 1.53 | 0.39 |
| Sensation Seeking | 2.85 | 0.70 |
| Delay Discounting Rate (K) | 0.02 | 0.06 |
| TLFB - Drinks Per Thirty Days | 20.19 | 13.69 |
| TLFB - Heavy Drinking Days | 19.08 | 10.96 |

**Notes.** Abbreviations as follow: BIS- Barratt Impulsiveness Scale; TLFB- Alcohol Timeline Followback.

### Measures

#### Alcohol Drinking Measures

The amount of daily alcohol consumption over the last 90 days was measured using the Alcohol Timeline-Followback (TLFB) calendar - a drinking assessment method with good psychometric characteristics for estimating retrospective daily drinking patterns [20]. The number of heavy drinking days in the last 90 days was calculated from the TLFB; a heavy drinking day is defined as 4 or more drinks per day for women and 5 or more drinks per day for men. We used this as a measure for binge drinking. Total drinks in the past 30 days was also calculated from the TLFB. Past 30-day drinking was used rather than 90-day drinking given that we were interested in recent drinking, and that recent reports of drinking may be more reliable.

#### Impulsivity Measures

Impulsive behavioral traits were measured using the Barratt Impulsiveness Scale (BIS-11) and the UPPS-P Impulsive Behavior Scale, two of the most widely-used self-report tools in the evaluation of trait impulsivity [21]. The BIS-11 scale evaluates impulsive behavior in general, and the motor, attentional and non-planning sub-scales evaluate acting without thinking, inability to focus attention and lack of forethought, respectively. The UPPS-P measures personality traits conducive to impulsive behavior including **U**rgency (negative) – tendency to impulsively act under strong negative emotions, **P**remeditation (lack of) – tendency to act without thinking, **P**erseverance (lack of) to remain focused on a task, **S**ensation-seeking – tendency to seek out novel/thrilling experiences, and **P**ositive-urgency - tendency to impulsively act under strong positive emotions.

We also evaluated choice impulsivity with the Delay Discounting Task (DDT). The DDT [22, 23] measures impulsivity by presenting the subject with a series of hypothetical choices between receiving a smaller immediate monetary reward or a larger delayed reward. The rate of discounting of the delayed outcome (k) was calculated and then a natural log-transformation was applied to correct for the non-normal distribution of k values. The resulting ln(k) is used to represent delay discounting, where higher ln(k) values mean greater preference for immediate rewards (see Table 1 for descriptive statistics of drinking and impulsivity scores for the sample).

### Experimental Design and Statistical Analysis

#### Overall Timeline

In this two-session study, healthy light drinkers received an IV alcohol infusion on two separate days. The first session was conducted outside the scanner to establish the alcohol infusion rate profile to achieve and maintain the target BrAC exposure, and to ensure tolerability. The alcohol infusion for the second session was conducted inside the scanner at least three days after the first session (Figure 1). Participants were asked to fast the night before each session. At the start of each visit, breath alcohol levels, urine pregnancy tests (if applicable) and urine drug screens were collected. In addition, we conducted a brief history interview about recent alcohol, medication, and nicotine use, changes in physical health, and menstruation (if applicable).

**Figure 1.**
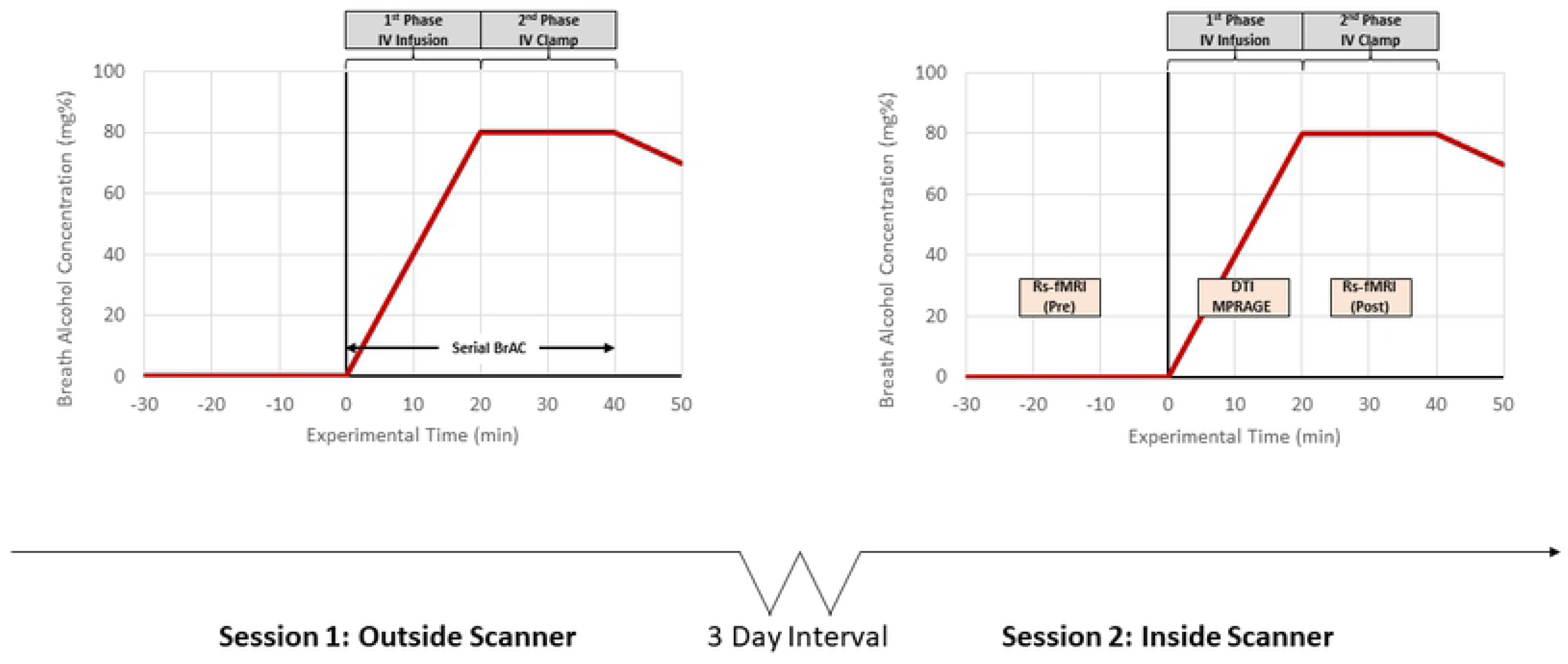
The timeline depiction of the two-session intravenous alcohol infusion study; sessions were separated at least three days from each other. The session 2 timeline indicates timepoints of resting state and BAC measurement alongside IV infusion; the first resting state scan occurred before the IV infusion at blood alcohol level of 0.0 g/dl and the second one was collected when the blood alcohol level reached the binge levels (0.08 g/dl).

#### Alcohol Infusion to Binge Level Exposure

We performed the alcohol infusion procedure following previously published methods (fMRI and PET) [24, 25]. In both sessions, participants received an intravenous (IV) infusion of 6% v/v ethanol solution in saline to achieve a target breath alcohol concentration (BrAC) of 0.08 ± 0.005 g/dl at 15 min after the start of the infusion, and to maintain (or clamp) the target BrAC level for 30 min (Figure 1). The infusion-rate profile was computed for each subject using a physiologically-based pharmacokinetic (PBPK) model-based algorithm, with individualized estimates of the model parameters estimated from the subject’s height, weight, age and sex [26, 27].

During the first session conducted outside the scanner, serial BrACs were measured at frequent intervals (5-15 minutes) using the Alcotest 7410+ handheld breathalyzer (Drager Safety Inc., CO), to ensure that each BrAC was within 0.005 g/dl of the target, and to enable minor adjustments to the infusion rates to overcome errors in parameter estimation and experimental variability [28, 29]. The adjusted infusion rate profile from the first session was used in the imaging session to replicate the target BrAC profile for each subject. This approach has been used to successfully achieve and maintain target BrACs, as verified by blood alcohol concentrations measured in samples drawn during the scan in other neuroimaging studies [25, 30]. After the end of the infusion, subjects were provided a meal, and BrAC was tracked until it dropped to 0.02 g/dL or below, at which time subjects were taken home by a designated driver or taxi.

#### Neuroimaging Acquisition and Preprocessing

During session 2, after the IV catheters were placed for alcohol infusion, participants were placed in a 3T SIEMENS Skyra MR scanner at the NMR Research Center at the NIH. Before starting the IV infusion of alcohol, a 5-min closed eyes resting-state fMRI scan was acquired with an echo-planar imaging sequence (36 axial slices, 3.8 mm thickness, 64 × 64 matrix and repetition time of 2000 ms, echo time 30, flip angle 90). After collecting this baseline resting-state fMRI scan, the alcohol infusion started and once the target BAC was achieved, a 10-minute waiting period was allowed to ensure the BAC was stable at the 0.08 g/dl level. During the waiting period, whole-brain structural and diffusion-weighted images of the brain were collected. Then, a second eyes-closed resting-state fMRI scan was collected while the target BAC (0.08 g/dl) was maintained. Following the MRI session, the Drug Effects Questionnaire was used to evaluate and quantify subjective effects of alcohol (DEQ) [31].

CONN ver.17f (http://www.conn-toolbox.org), a MATLAB/SPM-based (www.fil.ion.ucl.ac.uk/spm/) software, was used to conduct spatial/temporal preprocessing and analyses [32]. Spatial preprocessing included slice-timing correction, realignment, co-registration, normalization, and spatial smoothing (8 mm). Using the default settings in CONN, images were resliced into isotropic 2mm voxels. Anatomical volumes were segmented into grey matter, white matter, and cerebrospinal fluid (CSF) areas and the resulting masks were eroded to minimize partial volume effects. At the individual level, the temporal timeseries of the participant-specific six rotation-translation motion parameters and the timeseries from within the white matter/CSF masks were used as temporal covariates, and removed from the blood oxygen level-dependent (BOLD) functional data using linear regression. The resulting residual BOLD timeseries were band-pass filtered (0.008Hz < f < 0.15Hz) [33].

#### Neuroimaging Analysis

To our knowledge there is no reported sub-division of human GPe with concrete landmark definition and/or Talairach coordinate demarcation. Therefore, for these analyses, we used right and left GPe masks obtained from the Lead-DBS toolbox (http://www.lead-dbs.org/). These anatomical masks were generated using manual tracing of several structural MRIs and subsequent co-registration with histological studies, and have a combined volume of approximately 929 mm^3^ [34]. Functional connectivity measures were computed at the single-subject level as a Pearson’s correlation coefficient between the signal time-series averaged across the GPe seed mask and the signal time-series of all other voxels (seed-to-voxel analysis) of the brain. These correlations were calculated separately for the pre-infusion and post-infusion scans. Correlation coefficients were then converted to normally distributed z-scores using the Fisher transformation. Analyses for right and left GPe seeds were conducted separately. For each of the pre- and post-infusion analyses 150 time points (across 300 seconds) were used.

To examine group level effects, the resulting single-subject connectivity maps were used as input for a paired t-test where the within-subject effect was timepoint (i.e., post-versus pre-alcohol infusion; see supplemental materials for report of the main effects of alcohol administration); Supplemental Table 1 indicates clusters identified as showing significant changes from pre to post alcohol infusion, as well as associated statistics. Between-subject effects of impulsivity (BIS-11, UPPS-P, and DDT) and recent drinking behaviors (number of drinks in the last 30 days, number of heavy drinking days) on the within-subject effect of timepoint were examined using a general linear model. For the seed-to-voxel analysis, reported clusters survived a height threshold of uncorrected p < 0.005 with a cluster level extent threshold of FDR-corrected p < 0.05. In the figures, these clusters are shown overlaid on structural brain cutaways. Corresponding line plots are only for visual interpretation of these effects.

In order to test our hypothesis that the interaction between recent drinking behaviors and impulsivity would contribute to alcohol-induced changes in GPe functional connectivity, we conducted multiple regression analysis in R. We modeled drinking history, impulsivity, and the interaction between the two on the dependent variable of change in GPe connectivity. We conducted these analyses only for significant clusters identified in the seed-to-voxel analysis. For the drinking measures identified in that seed-to-voxel analysis, models (for the specific significant cluster only) included the significant drinking variable as well as one of the six impulsivity variables; thus, six regressions were run per cluster. Similarly, for significant impulsivity measures, 2 regressions per cluster were run (as there are two drinking variables). This resulted in a total of 110 tests. Reported results survived correction for multiple comparisons using FDR of < 0.05 at the omnibus test level. Of those regressions, only those with significant interaction effects are reported.

## Results

### GPe connectivity and Drinking Behaviors

Neither the total number of drinks over 30 days nor the number of heavy drinking days were correlated with subjective effects of the alcohol infusion as measured by the DEQ. Table 2 shows the correlations between drinking and impulsivity measures.

**Table 2.** Correlations between Drinking and Impulsivity Measures

| Measure | Delay Discounting Rate (K) | TLFB: Heavy Drinking Days | TLFB: Drinks Per Thirty Days | UPPS-P: Positive Urgency | UPPS-P: Negative Urgency | UPPS-P: Perseverance | BIS: Attentional Impulsiveness | BIS: Motor Impulsiveness | BIS: Nonplanning Impulsiveness |
| --- | --- | --- | --- | --- | --- | --- | --- | --- | --- |
| TLFB: Heavy Drinking Days | -0.22 | - | - | - | - | - | - | - | - |
| TLFB: Drinks Per Thirty Days | -0.33 | 0.79 *** | - | - | - | - | - | - | - |
| UPPS-P: Positive Urgency | 0.07 | 0.27 | 0.30 | - | - | - | - | - | - |
| UPPS-P: Negative Urgency | 0.22 | 0.30 | 0.14 | 0.48 * | - | - | - | - | - |
| UPPS-P: Perseverance | 0.06 | -0.28 | -0.22 | 0.06 | 0.46 * | - | - | - | - |
| BIS: Attentional Impulsiveness | 0.11 | 0.29 | 0.15 | 0.36 | 0.42 * | 0.13 | - | - | - |
| BIS: Motor Impulsiveness | 0.01 | 0.05 | 0.25 | 0.38 | 0.17 | 0.36 | 0.59 ** | - | - |
| BIS: Nonplanning Impulsiveness | 0.37 | -0.07 | -0.06 | 0.18 | 0.19 | 0.59 ** | 0.36 | 0.64 *** | - |
| BIS: Total | 0.22 | 0.09 | 0.11 | 0.36 | 0.30 | 0.46 * | 0.75 *** | 0.89 *** | 0.84 *** |

**Notes.** Abbreviations as follow: BIS- Barratt Impulsiveness Scale; TLFB- Alcohol Timeline Followback. Significance as follows- \*: $p < 0.05$ ; \*\*: $p < 0.01$ ; \*\*\*: $p < 0.001$

#### Number of drinks in 30 days

The number of drinks consumed per 30 days prior to screening had a significant effect on alcohol-related change in connectivity. Drinks in past 30-days was negatively associated with alcohol-related changes in the connectivity between the left GPe and the left hippocampus /amygdala, left IFG extending into the anterior insula, and areas of the bilateral superior frontal / paracingulate cortices. Drinks in past 30-days was also negatively associated with alcohol-induced change in connectivity between right GPe and bilateral putamen, Heschl’s gyrus, posterior insula, and central opercular cortices (Figures 2 and 5; Table 3). For each of these associations, the change was such that in individuals with more past 30 days drinking, alcohol decreased connectivity; in individuals with less drinking, alcohol increased connectivity.

**Table 3.** Associations between Alcohol Induced Changes in GPe Connectivity and Drinking Behaviors

|  |  | Peak |  |  | Cluster Size | p-FDR | p-FWE | Anatomical Labels |
| --- | --- | --- | --- | --- | --- | --- | --- | --- |
|  |  | x | y | z |  |  |  |  |
| DRINKING BEHAVIOR CORRELATIONS | Timeline Followback |  |  |  |  |  |  |  |
|  | Right GPe x 30 day drinking<br>(Negative correlation) | -42 | -16 | 6 | 100 | 0.023 | 0.014 | Left Heschl's Gyrus extending into the Insula, Central Operculum, Putamen |
|  |  | 42 | -20 | 10 | 83 | 0.029 | 0.036 | Right Heschl's Gyrus into the Insula, Planum Temporale, Central/Parietal Operculum, Putamen |
|  | Right GPe x 30 day drinking<br>(Positive correlation) | -12 | 18 | -10 | 98 | 0.008 | 0.016 | Left Subgenual Anterior Cingulate extending into the Accumbens |
|  |  | 6 | 18 | -12 | 75 | 0.015 | 0.057 | Right Subgenual Anterior Cingulate extending into the Accumbens |
|  | Left GPe x 30 day drinking<br>(Negative Correlation) | -2 | 50 | 20 | 142 | 0.002 | 0.001 | Superior Frontal Gyrus extending into dorsal anterior cingulate |
|  |  | -34 | 28 | 2 | 72 | 0.042 | 0.062 | Left Inferior Frontal Gyrus (pars triangularis) extending into Insula / Frontal Operculum |
|  |  | -42 | -8 | -26 | 66 | 0.042 | 0.089 | Left Hippocampus / Amygdala |
|  | Left GPe x Heavy drinking days<br>(Negative Correlation) | -38 | -8 | -28 | 291 | 0.002 | 0.003 | Left Parahippocampal Gyrus / Amygdala / Fusiform cortex |
|  | Left GPe x Heavy drinking days<br>(Positive correlation) | -18 | 18 | -14 | 215 | 0.017 | 0.020 | Left Orbitofrontal cortex extending into subgenual anterior cingulate and frontal medial cortex |
|  |  | -40 | -56 | 46 | 166 | 0.033 | 0.078 | Left Angular / Supramarginal Gyrus |
|  | Right GPe x Heavy drinking days<br>(Negative Correlation) | -36 | -22 | 6 | 279 | 0.005 | 0.005 | Left Insula extending into Heschl's Gyrus, Putamen, Pallidum, and Planum Temporale |
|  | Right GPe x Heavy drinking days<br>(Positive correlation) | -14 | 18 | -14 | 207 | 0.018 | 0.029 | Left Orbitofrontal cortex extending into subgenual anterior cingulate, frontal medial cortex, and accumbens |
|  |  | 6 | 68 | -4 | 189 | 0.018 | 0.046 | Bilateral Frontal Pole |

**Figure 2.**
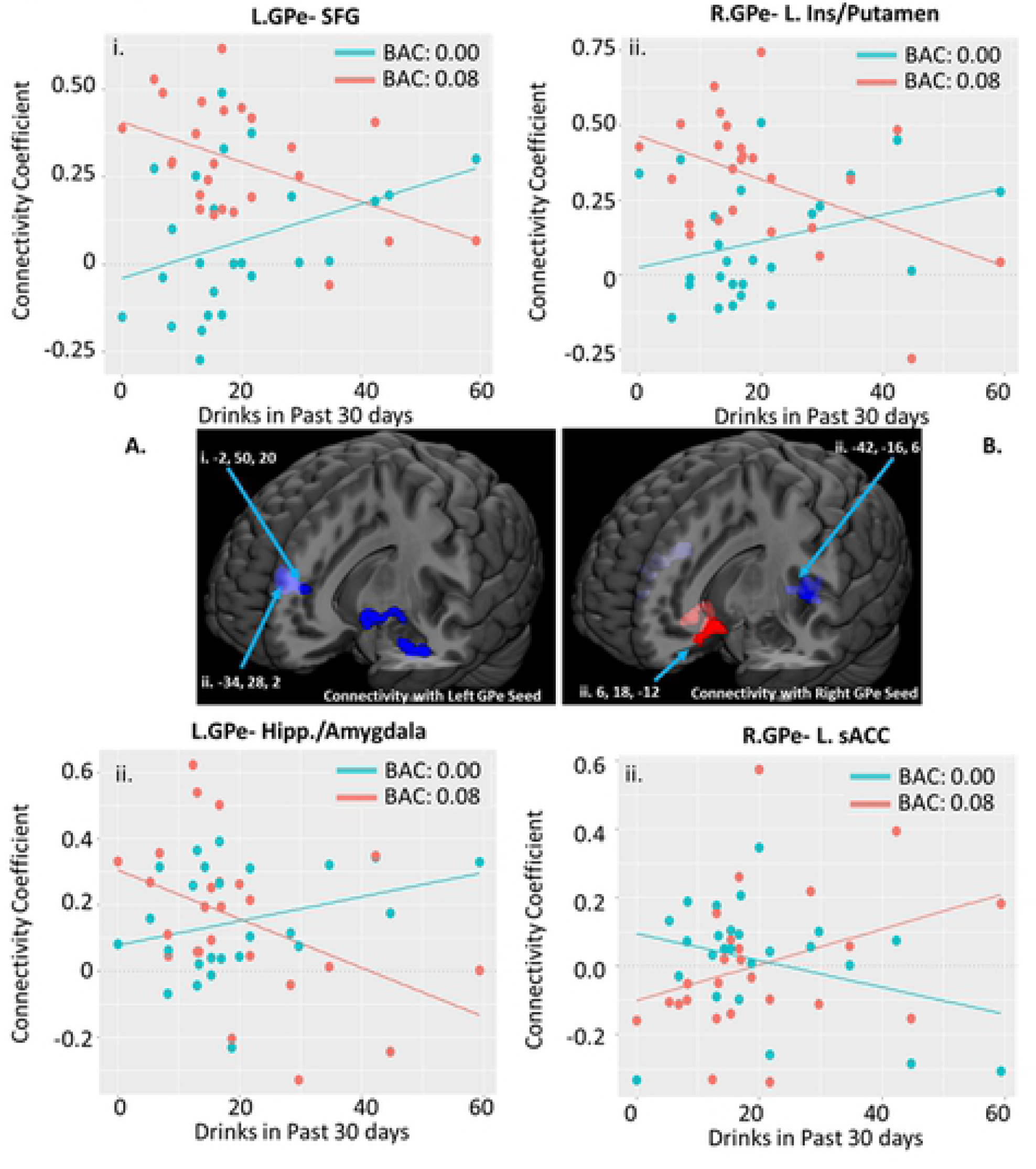
Association between Past 30 Day Drinking (as measured by the Alcohol Timeline Followback) and Alcohol Induced Changes in GPe Connectivity. Scatter plots correspond to individual connectivity at baseline in teal (estimated BAC: 0.0) and after infusion in salmon (BAC: 0.08); linear fit lines are also displayed for each timepoint. Connectivity coefficient is the r (correlation value) between the signal in the GPe seed and the coordinate indicated. Brain images represent whole effects of drinks in past 30 days on alcohol induced change in functional connectivity (pre-infusion connectivity > post-infusion connectivity). Warm colors indicate intoxication related increases in connectivity; cool colors indicated intoxication related decreases in connectivity. Images shown at p < 0.005. Clusters with significant interaction effects are reported in Figure 5. Reported clusters of connectivity with GPe area correspond to the following regions: A.i. – Superior Frontal Gyrus extending into dorsal anterior cingulate; A.ii. – Hippocampus/amygdala; B.i. – Heschl’s Gyrus extending into the insula/putamen; B.ii. - Heschl’s Gyrus extending into the insula/putamen.

**Figure 3.**
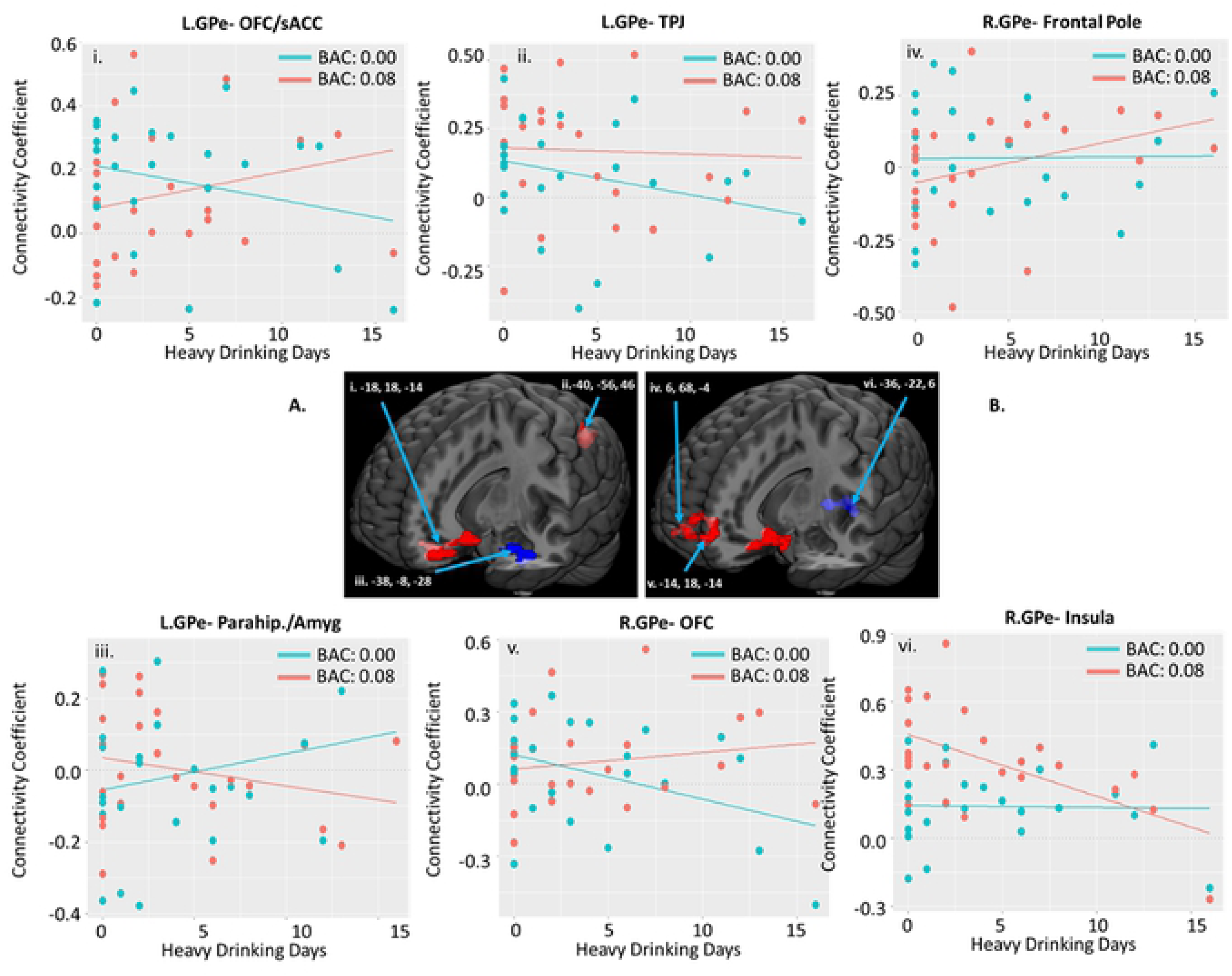
Association between Heavy Drinking Days (as measured by the Alcohol Timeline Followback) and Alcohol Induced Changes in GPe Connectivity. Scatter plots correspond to individual connectivity at baseline in teal (estimated BAC: 0.0) and after infusion in salmon (BAC: 0.08); linear fit lines are also displayed for each timepoint. Connectivity coefficient is the r (correlation value) between the signal in the GPe seed and the coordinate indicated. Brain images represent whole effects of drinks in past 30 days on alcohol induced change in functional connectivity (pre-infusion connectivity > post-infusion connectivity). Warm colors indicate intoxication related increases in connectivity; cool colors indicated intoxication related decreases in connectivity. Images shown at p < 0.005. Clusters with significant interaction effects are presented in Figure 5. Reported clusters clusters of connectivity with GPe area correspond to the following regions: i. – Orbitofrontal cortex extending into subgenual anterior cingulate; ii. – Angular/supramarginal gyrus; iii. Frontal pole.

**Figure 4.**
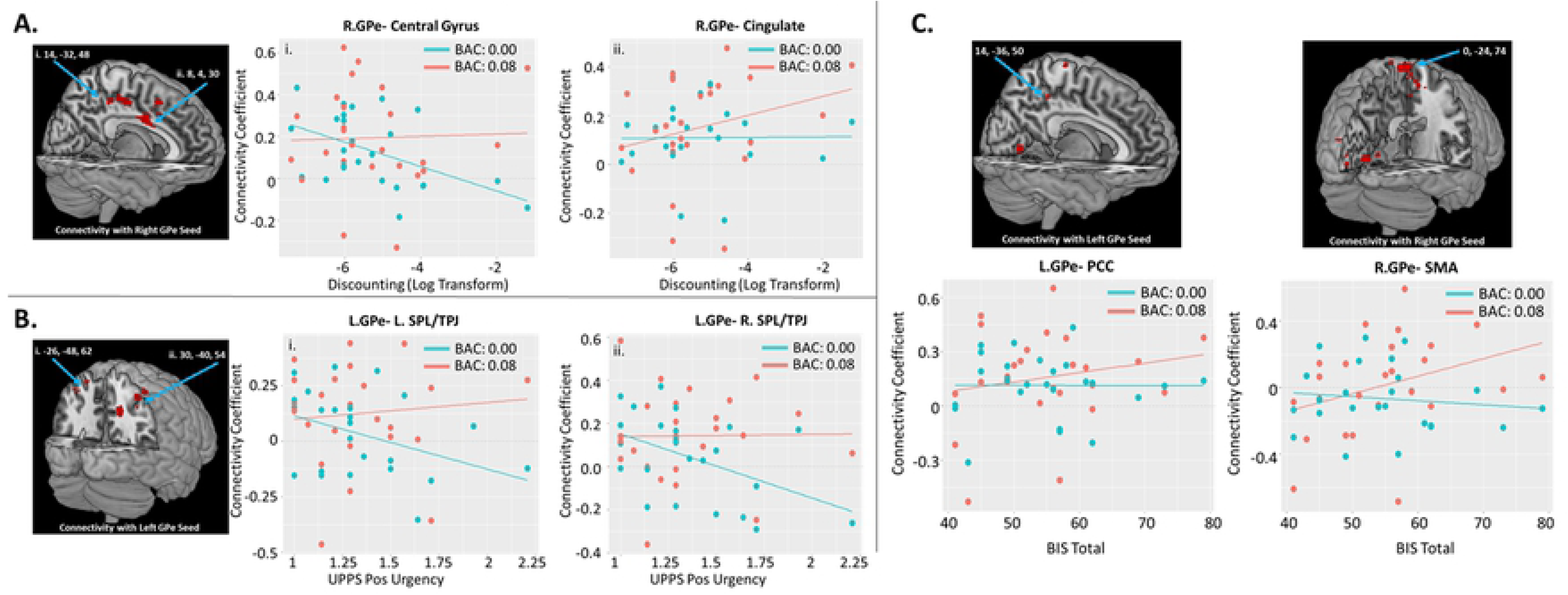
Association between Impulsivity Measures and Alcohol Induced Changes in GPe Connectivity. Scatter plots correspond to individual connectivity at baseline in teal (estimated BAC: 0.0) and after infusion in salmon (BAC: 0.08); linear fit lines are also displayed for each timepoint. Connectivity coefficient is the r (correlation value) between the signal in the GPe seed and the coordinate indicated. Brain images represent whole effects of drinks in past 30 days on alcohol induced change in functional connectivity (pre-infusion connectivity > post-infusion connectivity). Warm colors indicate intoxication related increases in connectivity. Images shown at p < 0.005. A) Changes in connectivity plotted by delay discounting task results. Values plotted are ln(k), where k is the discounting rate. Greater discounting rates indicate more choice impulsivity. B) Changes in connectivity plotted by scores on the positive urgency subscale of the UPPS-P impulsivity measure. C) Changes in connectivity plotted by total scores on the BIS impulsivity measure. Reported clusters of connectivity with GPe area correspond to the following regions: A.i. – Precentral/postcentral gyrus; A.ii. – Mid cingulate cortex extending anteriorly and posteriorly; B.i. – Superior parietal lobule extending into supramarginal gyrus; B.ii. – Superior parietal lobule extending into supramarginal gyrus; C.i. – postcentral gyrus extending into posterior cingulate; C.ii. – precentral gyrus/SMA

**Figure 5.**
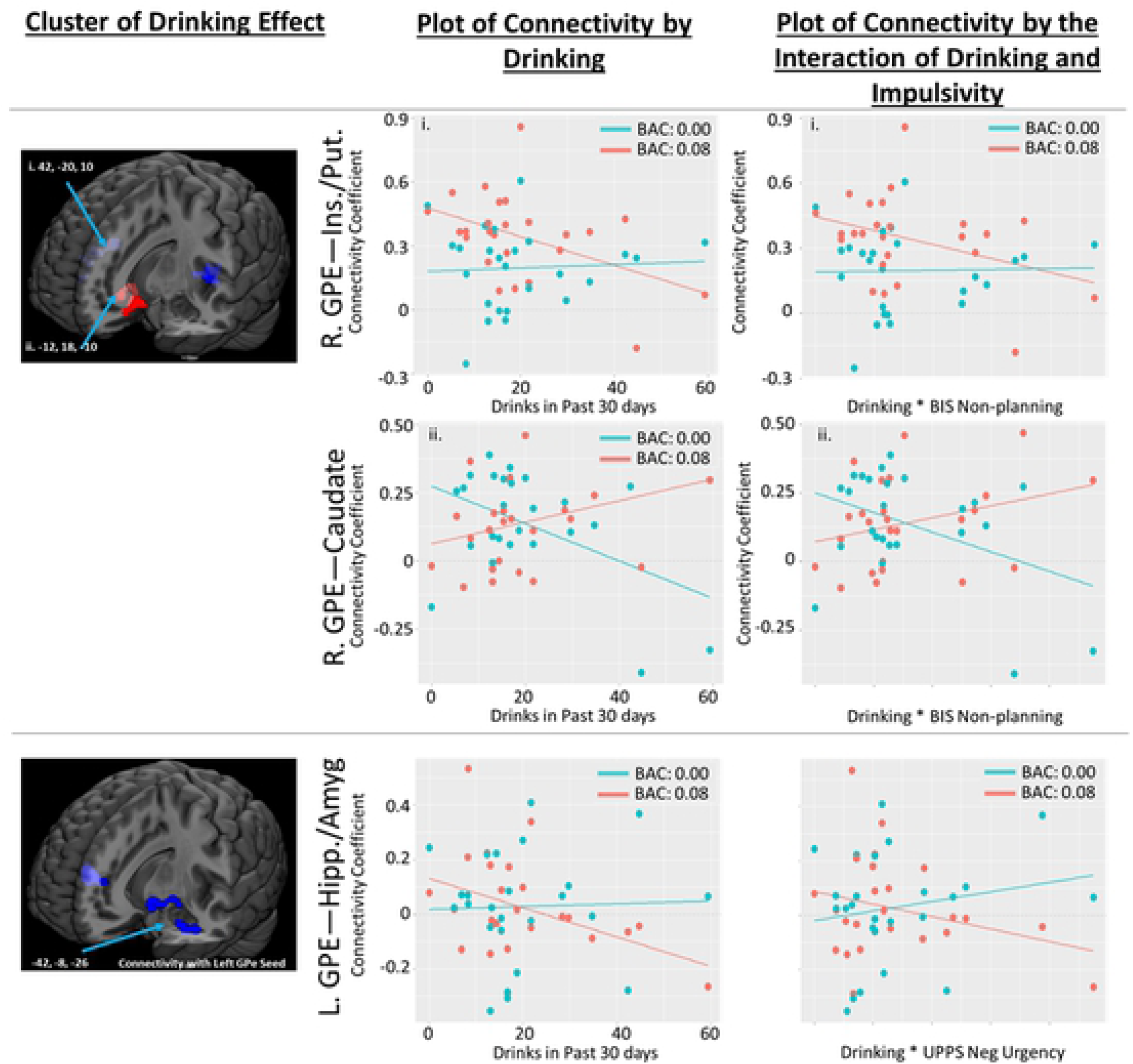
Effect of Alcohol Induced Changes in GPe Connectivity on the Association between Drinking (as measured by the Alcohol Timeline Followback) and Impulsivity (as measured by the BIS, UPPS, and delay discounting tasks). Cluster of Drinking Effect column: Brain images represent whole effects of drinks in past 30 days on alcohol induced change in functional connectivity (pre-infusion connectivity > post-infusion connectivity). Warm colors indicate intoxication related increases in connectivity. Images shown at p < 0.005. Plot of Connectivity by Drinking column: Scatter plots correspond to individual connectivity at baseline in teal (estimated BAC: 0.0) and after infusion in salmon (BAC: 0.08); linear fit lines are also displayed for each timepoint. Connectivity coefficient is the r (correlation value) between the signal in the GPe seed and the coordinate indicated. Diagram of Drinking/Impulsivity Mediation column: plot of mediation model testing. a = correlation between drinking and change in brain activity (post-pre); b = mediation effect, c = total effect, c’ = direct effect. Values indicate path estimates. Significance indicated as follows: p < 0.1t; p < 0.05*, p < 0.01**, p < 0.001***. Abbreviations as follows: GPe-Globus Pallidus externus; IFG-Inferior Frontal Gyrus; sACC-subgenual anterior cingulate; Parahip-Parahippocampal Gyrus; OFC-Orbitofrontal Cortex; L.-Left; R.-Right.

The number of drinks consumed in the past 30 days was positively associated with alcohol-induced changes in connectivity between the right GPe and the bilateral subgenual cingulate extending into the basal ganglia. The change was such that in individuals with more past 30 days drinking, alcohol increased connectivity; in individuals with less drinking, alcohol decreased connectivity.

#### Total number of Heavy Drinking Days

The total number of heavy drinking days was associated negatively with changes in left GPe ipsilateral connectivity with the fusiform cortex, hippocampus, amygdala and parahippocampal cortex, such that individuals with more heavy drinking showed an alcohol-related decrease in connectivity. A negative association with number of heavy drinking days was also found in connectivity between the right GPe and left areas of the IFG, Heschl’s gyrus, insular cortex, putamen, pallidum and planum temporale, such that there were no drinking pattern related differences in connectivity at baseline; however, in individuals with fewer heavy drinking days but not those with more, alcohol infusion increased connectivity (Figures 3 & 5; Table 2).

Number of heavy drinking days was positively associated with connectivity changes following alcohol infusion between the left GPe and left frontal pole extending into medial prefrontal cortex (mPFC), bilateral subgenual ACC, OFC, and angular gyrus extending into the superior division of left lateral occipital cortex. We also found positive associations between the number of heavy drinking days and connectivity between right GPe and the NAcc, as well as ACC and frontal (mPFC/OFC, frontal pole) regions. This pattern of alcohol-related connectivity was such that individuals with more heavy drinking days had less connectivity at baseline with these clusters; alcohol also increased connectivity between GPe and the OFC clusters.

##### GPe connectivity and Impulsivity

Measures of impulsivity (BIS-11, UPPS-P, DDT) were not correlated with subjective effects of alcohol infusion as measured by the DEQ. Although we did examine both positive and negative associations, there were no significant negative associations between alcohol-induced GPe connectivity changes and impulsivity. Other findings reported below.

#### BIS-11 Trait Impulsivity

We found a significant positive association between BIS-11 total score and alcohol - induced connectivity change of left GPe with areas of the bilateral postcentral and posterior cingulate gyri. Change in connectivity between the right GPe with the precentral gyrus also showed significant positive association with the BIS-11 total score. In other words, higher impulsivity score was associated with increased GPe functional connectivity while under the influence of alcohol; there was no association between impulsivity and connectivity with those clusters at baseline. Further examination of the BIS-11 indicated that the GPe connectivity in these areas were positively associated with the BIS-11 non-planning sub-scores, which is related to impulsive behaviors due to lack of forethought. BIS-11 non-planning score was also positively correlated with the alcohol-induced GPe connectivity changes in the bilateral cuneus/precuneus, posterior cingulate gyri, supra and intracalcarine cortices, superior division of the lateral occipital cortices, occipital pole and left lingual gyrus (Figures 4A, 4B and Table 4). Other BIS sub-scores were not significantly related to change in connectivity.

**Table 4.**
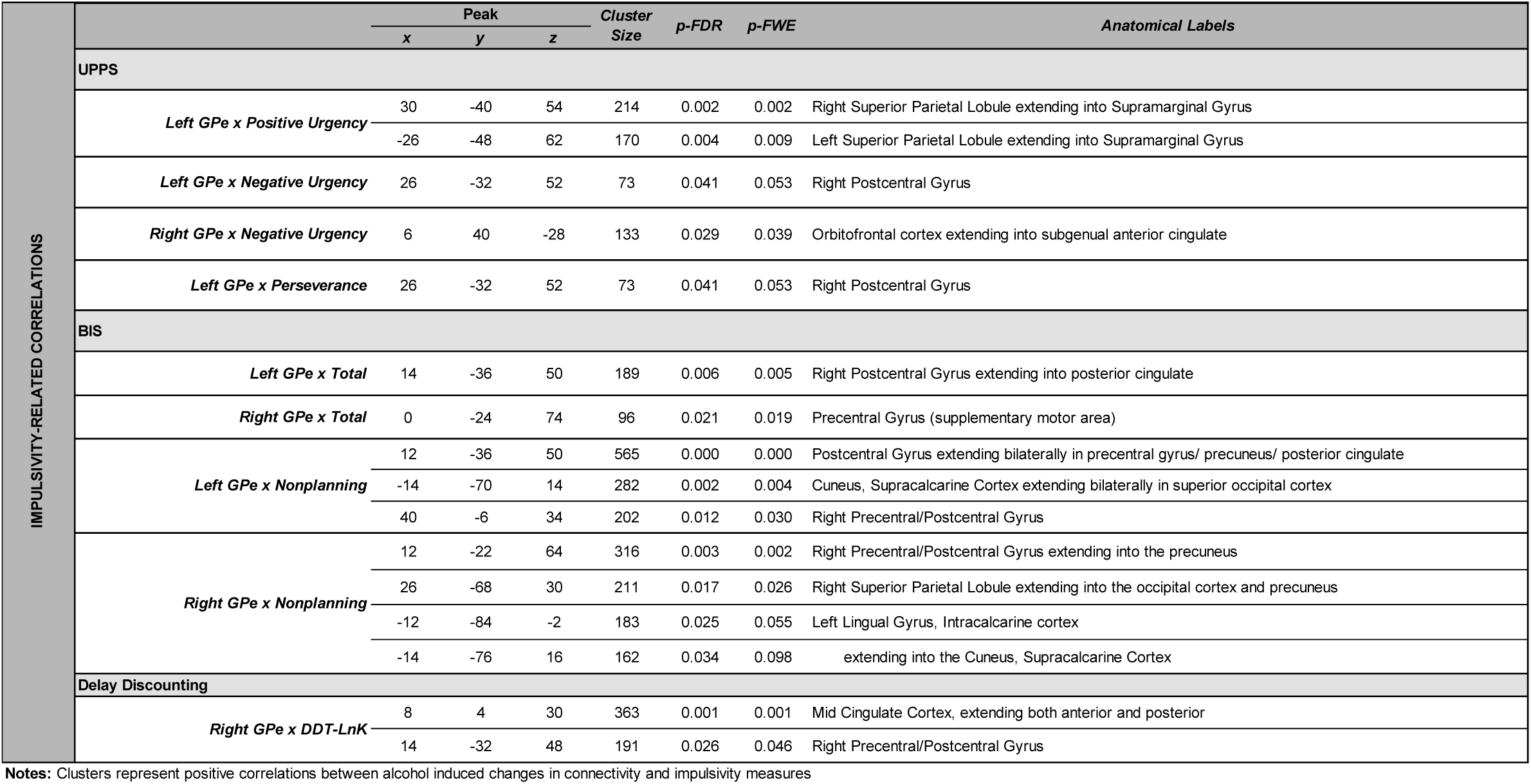
Associations between Alcohol Induced Changes in GPe Connectivity and Impulsivity Measures

|  |  | Peak |  |  | Cluster Size | p-FDR | p-FWE | Anatomical Labels |
| --- | --- | --- | --- | --- | --- | --- | --- | --- |
|  |  | x | y | z |  |  |  |  |
| IMPULSIVITY-RELATED CORRELATIONS | UPPS |  |  |  |  |  |  |  |
|  | Left GPe x Positive Urgency | 30 | -40 | 54 | 214 | 0.002 | 0.002 | Right Superior Parietal Lobule extending into Supramarginal Gyrus |
|  |  | -26 | -48 | 62 | 170 | 0.004 | 0.009 | Left Superior Parietal Lobule extending into Supramarginal Gyrus |
|  | Left GPe x Negative Urgency | 26 | -32 | 52 | 73 | 0.041 | 0.053 | Right Postcentral Gyrus |
|  | Right GPe x Negative Urgency | 6 | 40 | -28 | 133 | 0.029 | 0.039 | Orbitofrontal cortex extending into subgenual anterior cingulate |
|  | Left GPe x Perseverance | 26 | -32 | 52 | 73 | 0.041 | 0.053 | Right Postcentral Gyrus |
|  | BIS |  |  |  |  |  |  |  |
|  | Left GPe x Total | 14 | -36 | 50 | 189 | 0.006 | 0.005 | Right Postcentral Gyrus extending into posterior cingulate |
|  | Right GPe x Total | 0 | -24 | 74 | 96 | 0.021 | 0.019 | Precentral Gyrus (supplementary motor area) |
|  | Left GPe x Nonplanning | 12 | -36 | 50 | 565 | 0.000 | 0.000 | Postcentral Gyrus extending bilaterally in precentral gyrus/ precuneus/ posterior cingulate |
|  |  | -14 | -70 | 14 | 282 | 0.002 | 0.004 | Cuneus, Supracalcarine Cortex extending bilaterally in superior occipital cortex |
|  |  | 40 | -6 | 34 | 202 | 0.012 | 0.030 | Right Precentral/Postcentral Gyrus |
|  | Right GPe x Nonplanning | 12 | -22 | 64 | 316 | 0.003 | 0.002 | Right Precentral/Postcentral Gyrus extending into the precuneus |
|  |  | 26 | -68 | 30 | 211 | 0.017 | 0.026 | Right Superior Parietal Lobule extending into the occipital cortex and precuneus |
|  |  | -12 | -84 | -2 | 183 | 0.025 | 0.055 | Left Lingual Gyrus, Intracalcarine cortex |
|  |  | -14 | -76 | 16 | 162 | 0.034 | 0.098 | extending into the Cuneus, Supracalcarine Cortex |
|  | Delay Discounting |  |  |  |  |  |  |  |
|  | Right GPe x DDT-LnK | 8 | 4 | 30 | 363 | 0.001 | 0.001 | Mid Cingulate Cortex, extending both anterior and posterior |
|  |  | 14 | -32 | 48 | 191 | 0.026 | 0.046 | Right Precentral/Postcentral Gyrus |

Notes: Clusters represent positive correlations between alcohol induced changes in connectivity and impulsivity measures

#### UPPS-P Trait Impulsivity

Tendency to display impulsive behaviors under strong positive emotions, as measured by the UPPS-P positive urgency score, was significantly positively associated with alcohol-induced connectivity changes between the left GPe and bilateral areas of the superior parietal lobules (SPL), supramarginal and postcentral gyri, such that baseline anticorrelated activity in individuals with higher UPPS-P positive urgency scores was normalized when intoxicated. The connectivity between the left GPe and right postcentral gyri was positively associated with the perseverance score of the UPPS-P, such that more impulsive individuals had greater connectivity both pre and post alcohol infusion. Alcohol-induced changes in right GPe connectivity with the postcentral, OFC, and the paracingulate gyri were positively associated with the UPPS-P negative urgency score (Figure 4C; Table 4). Individuals with higher negative urgency scores had lower connectivity with these clusters at baseline. In addition, the OFC cluster had increased alcohol related connectivity associated with impulsivity.

#### DDT Ln(k) Choice Impulsivity

The delay discounting ln(k) score of the DDT was positively associated with connectivity between the right GPe and areas of the precentral/post central gyrus and ACC/posterior cingulate cortices (Figure 4A; Table 4). Connectivity was not related to discounting at baseline with the cingulate, but individuals with higher discounting rates had greater connectivity while intoxicated. On the other hand, those individuals had lower connectivity at baseline with the pre/postcentral gyrus, but this normalized with alcohol infusion.

### Interaction between drinking and impulsivity on GPe connectivity change

The interaction between number of drinks in the past 30 days and scores on the BIS non-planning subscales significantly contributed towards alcohol induced change in right GPe connectivity with right Heschl’s gyrus/insula/putamen and caudate regions. Specifically, individuals with higher trait non-planning impulsivity scores and more recent drinking had no hyperconnectivity of the insula and GPe regions when intoxicated, while those low on both measures did. On the other hand, individuals high in both measures had little baseline connectivity between the GPe and caudate but notable connectivity between those regions when intoxicated; individuals low on both traits had the reverse pattern of alcohol effects where connectivity was observed at baseline, but not while intoxicated. The interaction between number of drinks in the past 30 days and UPPS negative urgency subscale scores also predicted alcohol-induced change between left GPe and the hippocampus / amygdala. Specifically, individuals with greater negative urgency and more recent drinking had greater connectivity between these regions at baseline, but anticorrelations between these regions when intoxicated. Individuals low on both of these measures had little connectivity in either conditions. See Table 5 for estimates and statistics; See Figure 5 for a graphical representation of these effects.

**Table 5.**
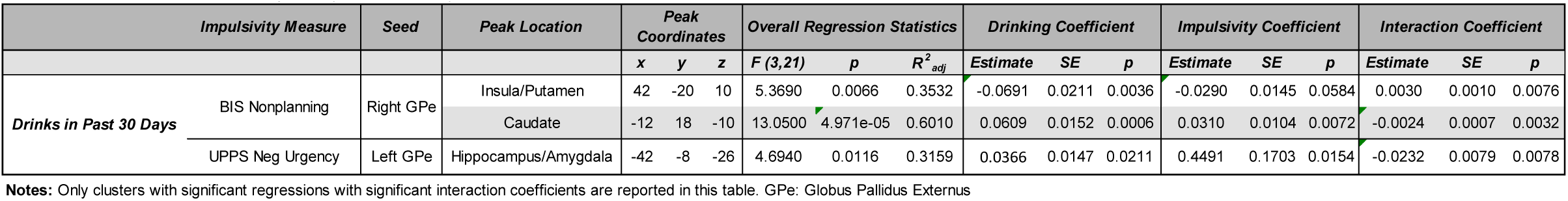
Interaction Between Drinking History and Impulsivity

## Discussion

In this study, we examined the association between changes in GPe resting state connectivity following alcohol infusion with drinking history and impulsivity. We hypothesized that alcohol infusion would more greatly reduce GPe functional connectivity in individuals with heavier drinking and more impulsivity. We also hypothesized that the interaction between recent drinking behaviors and impulsivity would contribute to alcohol-induced changes in GPe functional connectivity. We found reductions in connectivity between the GPe and some subregions of the brain following alcohol infusion in individuals with more drinking in the past 30 days and more heavy drinking days. However, impulsivity was related to greater GPe connectivity while intoxicated. Heavier recent drinking behaviors in combination with higher impulsivity were associated with greater alcohol-induced connectivity change between GPe and auditory, caudate, and amygdala regions, but for most of our connectivity findings, the contribution of drinking and impulsivity were distinct. Together, our findings suggest that the relationship between GPe-related inhibition and alcohol-related impulsivity is more complicated than we previously anticipated.

As predicted, we found that heavier drinking individuals had more pronounced changes in GPe connectivity with the basal ganglia, OFC, and mPFC regions. In these regions, individuals with patterns of heavy drinking had less coupling at baseline than individuals with lighter drinking patterns; in the binge infusion state, this association between heavy drinking history and GPe connectivity disappears, or in the case of the OFC, reverses. The pronounced baseline reduction of downstream GPe-frontal pathways in heavier drinkers may specifically reflect impairment of stop signaling [35]. Not only does this reduced behavioral inhibition correspond to a greater likelihood of binge drinking, the larger difference between states may reflect an alcohol sensitization effect, consistent with sensitization effects seen for other drugs of abuse, particularly in basal ganglia regions [36, 37].

We also found that heavier drinking individuals had less change caused by alcohol infusion in connectivity between the GPe and lateral IFG, ACC, putamen, insula, and amygdala/hippocampus regions. In fact, while alcohol decreased GPe connectivity with these regions in individuals with more recent drinking, the effect was reverse for those with less drinking. In the IFG, ACC, and amygdala/hippocampus, this may correspond to tighter coupling at baseline. The IFG has been demonstrated to control stop-signaling through the supplementary motor area, upstream from sub-cortical processes [38]. It may be that this increased connectivity in the IFG network reflects a shift towards reactive rather than proactive inhibition for individuals with more drinking [39].

Reward-driven overrides of the stop-signal from the dorsal ACC likely exacerbate differences in frontal-subcortical pathways [40]. Greater coupling of dACC-GPe at baseline may lead to a greater tendency to override the stop-signal in face of drinking; this would likely be more pronounced in individuals with more recent, positive experiences with alcohol. However, this override would be less necessary when the impulse control pathways are working sub-optimally and sending less stop-signals, such as during intoxication. On the other hand, in individuals with less drinking, alcohol may serve to reduce inhibition via increased connectivity to this override, explaining the differential pattern of results.

The insula is thought to play a role in signaling unsuccessful stopping, particularly in cases where errors are less common [41]. Increased intoxication-related changes in connectivity between this region and the GPe in individuals with less recent drinking, particularly those with low levels of trait impulsivity, may reflect novelty of failures in stop-signaling. Individuals who drink more may expect failures of stop-signaling during intoxication, as opposed to those with infrequent drinking. A lack of baseline association between recent drinks and error feedback in the right insula is consistent with lack of differences in expectation of low-frequency errors at baseline. Alternatively, during intoxication, less stop-signaling could correspond to less failures, leading to a reduced need for error signaling.

However, it is difficult to determine if lower baseline GPe connectivity is caused by heavy drinking or if it is a vulnerability factor that leads to heavy drinking. It may be that individuals with lower GPe-frontal connectivity (impaired stop-signaling) are more vulnerable to engaging in heavy drinking episodes. Another possibility is that drinking, even at a social level, has lasting effects on resting thresholds of GPe-frontal connectivity; some work has shown that alcohol leads to hyperactive IFG response during stop-signaling [18]. Future work should use longitudinal approaches to investigate risk factors and consequences of heavy and binge drinking across the lifespan.

Impulsivity across measures was associated with greater change in GPe connectivity following alcohol infusion. For clusters in the bilateral SPL/supramarginal gyrus, postcentral/precentral gyrus, and cingulate, individuals low on impulsivity had no differences (or in the case of the OFC, decreases) in connectivity with the GPe as a result of alcohol. In fact, across the continuum of impulsive traits, there were no baseline differences in GPe connectivity for clusters in the cingulate, precuneus, and lingual gyrus. However, after alcohol infusion, impulsivity was related to increased connectivity of these areas to the GPe. Previous work has shown that activation of this network of regions is associated with more errors in stop-signaling [42]. Alcohol may increase the resting activation of this network, leading to a larger tendency for failures of behavioral inhibition. This alcohol-induced impulsivity may be most pronounced in individuals who have greater impulsivity at a trait level.

Our results also suggest a baseline vulnerability to impulsivity in individuals who drink to cope with stress. We found hyperactivity in the amygdala-GPe circuit at baseline in individuals with greater impulsivity during negative emotionality who also have more recent drinking. Increased connectivity between the amygdala and basal ganglia is associated with greater stress-induced cortisol response dependent on the availability of mineralocorticoid receptors [43]. Our observation of decoupling during intoxication in individuals with high UPPS-P negative urgency suggests that individuals with a tendency towards impulsive behaviors when stressed may use alcohol as a mechanism to reduce stress response in this pathway. This “drinking to reduce negative affect” is a key feature in addiction [44], and may mean that individuals with this pattern of response are particularly vulnerable to developing alcohol use disorder. In fact, chronic alcohol consumption leads to increased mineralocorticoid receptor pathway activity [45], suggesting that by drinking to reduce stress, these individuals may be making themselves more vulnerable to stress in the future.

Alcohol-induced increases in connectivity between the GPe and other regions were associated with differences in baseline connectivity, including GPe-SPL decoupling in individuals with a tendency to be impulsive during positive emotionality, GPe-caudate decoupling in individuals with non-planning impulsivity as well as with more recent drinking, and low baseline GPe-precentral gyrus connectivity in individuals who devalue future rewards. Previous studies have shown that stop-signaling is associated with connectivity in frontoparietal/striatal (caudate extending into GPe) networks [46]. Moreover, other work has shown that parietal-striatal network engagement is indicative of proactive stop-signaling and negatively correlated with motor urgency [47], while precentral/SMA engagement is associated with reactive inhibition [39] and influences inhibition through a hyperdirect rather than indirect pathway [48]. Although these impulsive individuals show less connectivity in the indirect stop-signaling pathway, it may be that for those with a pre-existing hypoconnectivity, alcohol leads to a higher likelihood of recruitment of alternative pathways to compensate for alcohol related impairment. However, the highest instance of successful inhibition is when the hyperdirect and indirect pathways work in conjunction [12], meaning individuals with disproportional recruitment of the hyperdirect pathway may still have increased impulsivity.

### Limitations

Because we did not have a placebo infusion condition, we cannot rule out that pre- to post-infusion changes were not due to dynamic fluctuations within resting states. However, we do not believe this is the case. A previous study using an alcohol infusion paradigm with a full, double-blind crossover design found alcohol effects on resting-state connectivity compared to placebo [49]. Although that study examined network level connectivity changes rather than seed-based changes in subcortical regions as reported here, the fact that acute alcohol intoxication impacts connectivity during resting state has already been established. Moreover, this study is based on animal literature that found alcohol-induced changes in this pathway that are dose-dependent [17].

Our measures of impulsivity and drinking were primarily retrospective and obtained via self-report. Although we have designed our analysis to provide insight on the relationship between impulsivity, drinking, and alcohol-related changes in brain connectivity, we cannot comment on the directionality or causality of these relationships. Nonetheless, we did find evidence that connectivity during our laboratory “binge drinking” state was influenced by previous drinking and impulsive traits. Future studies should take a prospective approach that could speak directly to the neural mechanisms leading to problem drinking behaviors, as well as measuring within-subject changes in choice impulsivity during acute intoxication.

Another limitation to our inferences about baseline impulsivity and drinking measures is that our participants were light social drinkers with a limited range of alcohol use and an absence of AUD. This may explain why impulsivity and drinking measures were not significantly correlated in our sample, despite their clear link in the literature (e.g., [50]). Overall, our findings are most applicable to non-AUD binge drinking groups, such as adolescents and emerging adults, rather than in chronic heavy binge drinkers with AUD.

The small sample size of this study (n = 25) allowed us to test our specific hypotheses but may limit our ability to rule out the involvement of GPe connectivity to areas that were not found to be significant in this study. We limited our investigation to the connectivity of the hypothesized region of interest (GPe) and did not investigate other potential pathways of behavior control. Importantly, we do not assert that GPe pathways are the only mechanisms of control influenced by acute or chronic alcohol consumption or related to impulsivity. In fact, increased connectivity in behavioral control circuits suggests that alternative mechanisms or alternative recruitment of pathways are involved in failures of inhibition. Although outside the scope of the current study, future work should investigate those pathways during acute alcohol intoxication in a larger sample, and during specific stop-signaling tasks.

There are other potential explanations for our findings. First, given that our study looked at resting-state connectivity differences, rather than actual differences in stop-signaling, it is possible that observed changes are not directly related to behavioral control. Abnormal frontostriatal connectivity has also been implicated in anhedonia and psychomotor retardation [51], mental time-keeping [52], cognitive load during working memory [53], set-shifting [54], and long-term reward learning [55]. Changes in connectivity could also be driven by alcohol-related changes in neurotransmitter availability across the whole brain that may or may not translate directly to functional change; dopamine has been shown to modulate both resting and set-shifting based functional connectivity [56].

## Conclusion

Here we translated preclinical research to investigate how alcohol-induced GPe connectivity disruptions explain impulsive behaviors and drinking patterns in social drinkers. We found that impulsivity and alcohol use are associated at rest with impairment of GPe circuits implicated in typical stop-signaling processing. To our knowledge, this is the first demonstration of an association between impulsivity, drinking measures, and GPe connectivity changes under clamped alcohol exposure in humans. These findings suggest that individuals with impulsive traits and past drinking are particularly vulnerable to the control impairments of alcohol intoxication. Moreover, they have important implications with regard to identifying those most at risk for drinking-related consequences; they suggest a mechanism for how alcohol leads to impulsive behaviors, including excessive drinking. Finally, the neuronal connectivity changes induced by alcohol may represent a target for interventions that would mitigate the impact of binge drinking or AUD by reducing impulsivity. Although additional work, including longitudinal studies, are needed before these findings can be useful in a clinical setting, this study achieves the important first step of translating preclinical models of binge drinking to the human brain and behavior.

